# Genomics and telemetry suggest a role for migration harshness in determining overwintering habitat choice, but not gene flow, in anadromous Arctic Char

**DOI:** 10.1101/138545

**Authors:** Jean-Sébastien Moore, Les N. Harris, Jérémy Le Luyer, Ben J. G. Sutherland, Quentin Rougemont, Ross F. Tallman, Aaron T. Fisk, Louis Bernatchez

## Abstract

Migration is a ubiquitous life history trait with profound evolutionary and ecological consequences. Recent developments in telemetry and genomics, when combined, can bring significant insights on the migratory ecology of non-model organisms in the wild. Here, we used this integrative approach to document dispersal, gene flow and potential for local adaptation in anadromous Arctic Char from six rivers in the Canadian Arctic. Acoustic telemetry data from 124 tracked individuals indicated asymmetric dispersal, with a large proportion of fish (72%) tagged in three different rivers migrating up the same short river in the fall. Population genomics data from 6,136 SNP markers revealed weak, albeit significant, population differentiation (average pairwise *F*_ST_ = 0.011) and asymmetric dispersal was also revealed by population assignments. Approximate Bayesian Computation simulations suggested the presence of asymmetric gene flow, although in the opposite direction to that observed from the telemetry data, suggesting that dispersal does not necessarily lead to gene flow. These observations suggested that Arctic Char home to their natal river to spawn, but may overwinter in rivers with the shortest migratory route to minimize the costs of migration in non-breeding years. Genome scans and genetic-environment associations identified 90 outlier markers putatively under selection, 23 of which were in or near a gene. Of these, at least four were involved in muscle and cardiac function, consistent with the hypothesis that migratory harshness could drive local adaptation. Our study illustrates the power of integrating genomics and telemetry to study migrations in non-model organisms in logistically challenging environments such as the Arctic.

## Introduction

Migrations are a common feature of the life history of many animal species (Dingle 2014). Despite their obvious evolutionary and ecological consequences, migrations have been a challenging topic of study. Quantifying migratory phenotypes is difficult because they can occur over vast distances. Furthermore, while some migratory traits are amenable to laboratory studies and more classical quantitative genetic approaches (e.g. Berthold & Querner 1981; Roff & Fairbairn 1991), many migratory traits are impossible to recreate in experimental settings. Recent technological developments in telemetry techniques both on land (Kays *et al.* 2015) and underwater (Hussey *et al*. 2015) now allow observations of animal movement in the wild over broad spatial scales and with unprecedented levels of detail. Parallel to these developments is the exponential increase in the availability of genomic technologies for non-model organisms (Davey *et al.* 2011, Andrews *et al.* 2016). These new genomic tools make genome-wide assessments of genetic variation possible, thus offering novel ways to link genotypes to migratory phenotypes in the wild for non-model organisms (Liedvogel *et al.* 2011; Hess et al. 2014; Shafer *et al.* 2016; Franchini *et al.* 2017). The integration of telemetry and genomic datasets, therefore, provides a powerful approach to document the population level consequences of migrations and to better understand the genetic basis of migratory traits (Shafer *et al.* 2016).

One population level consequence of migration is that it can redistribute genetic variation. Indeed, while homing to reproduction sites is commonly associated with migrations (Dingle 2014), it is rarely perfect, thus leading to dispersal and gene flow. Consequently, the precision of homing and the spatial scale over which migrations take place influence the scale over which gene flow, and thus genetic structure, can be observed (e.g., Castric & Bernatchez 2004). Furthermore, inter-individual differences in migratory behaviour are common and can influence gene flow (e.g., Turgeon *et al.* 2012; Shafer *et al*. 2012; Delmore *et al*. 2015). Therefore, the study of how migratory behaviour, dispersal, and gene flow interact to determine the genetic structure of populations and their capacity to locally adapt can greatly benefit from integrating genomic and telemetry data.

The diversity of migratory life histories in salmonids makes them excellent model systems for the study of migration ecology (Hendry *et al*. 2004; Quinn 2005). Anadromy, a trait shared by many salmonid species, refers to a migratory life cycle whereby individuals are born in freshwater, feed and grow in saltwater, and return to freshwater to reproduce. In most salmonids, return migrations to freshwater occur exclusively for the purpose of reproduction, but some species, notably in the genus *Salvelinus*, must also return to freshwater to overwinter (e.g., Johnson 1980; Moore *et al.* 2013; Bond *et al*. 2015). The Arctic Char (*Salvelinus alpinus*) is a facultatively anadromous salmonid with a circumboreal distribution (Johnson 1980; Klemetsen *et al*. 2003; Reist *et al*. 2013). Anadromous individuals undergo annual feeding migrations to the ocean and must return to freshwater in the fall because winter conditions in the Arctic Ocean are not favorable (Johnson 1980; Klemetsen *et al*. 2003; see Jensen & Rikardsen 2012 for an exception). The summer feeding and growth period is thus limited by the ice-free period on the rivers used for migrations, which can be as short as a month at higher latitudes (Johnson 1980). This restricted feeding period limits energy gains, and results in skipped reproduction such that in most populations, spawning occurs once every two to four years (Dutil 1986). The rate of iteroparity is among the highest reported for anadromous salmonids, with 32-50% of individuals observed breeding more than once in some populations (Fleming 1998). The migratory behaviour of anadromous Arctic Char is therefore best understood as three distinct migrations: (1) spring feeding migrations to the ocean; (2) fall spawning migrations to freshwater spawning sites in headwater lakes; and (3) fall overwintering migrations to freshwater overwintering sites in headwater lakes when the individual is not in breeding condition (Fig. 1). Given that optimal spawning habitats likely differ from optimal overwintering habitats, we can predict that individuals might select different habitats in different years depending on their maturity status. Accordingly, there is evidence that Arctic Char home to their natal sites to spawn, but often utilize non-natal overwintering sites in years when they are not in breeding condition (Johnson 1980; Gyselman 1994; Moore *et al*. 2013; Gilbert *et al*. 2016). Natal homing to reproduction sites combined with the use of alternative overwintering sites may lead to temporary mixing of different stocks in freshwater, making both population-specific sampling and fisheries management challenging. The integration of telemetry and genomic datasets is therefore particularly promising for studying Arctic Char migrations.

**Figure 1.**
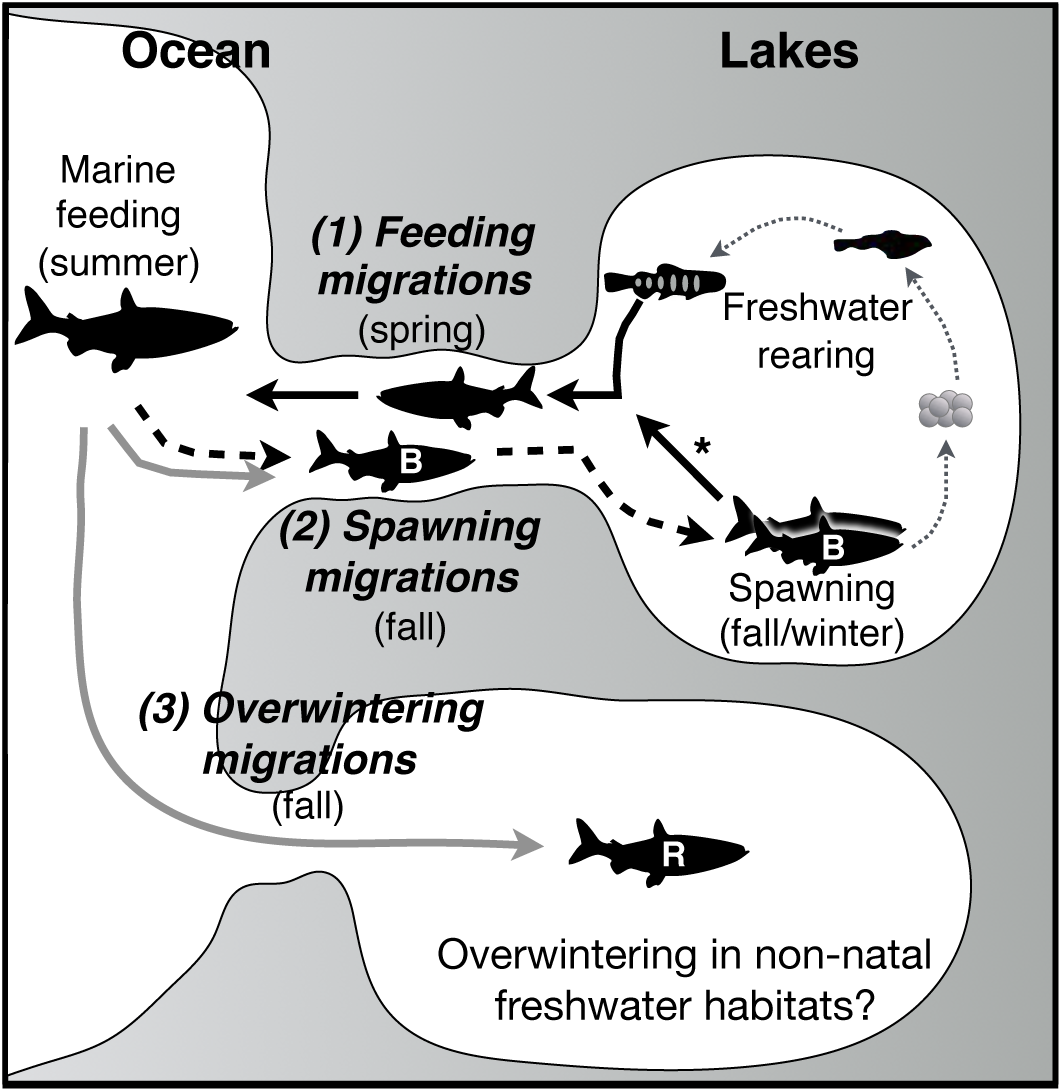
The anadromous Arctic Char life cycle highlighting three distinct migrations. (1) Feeding migrations (black arrows): after hatching and rearing in freshwater for several years (grey dotted arrows), individuals smoltify and begin to migrate to saltwater to feed during the summer when the rivers and ocean are free of ice. (2) Spawning migrations (dashed black arrows): individuals in breeding condition (B) return to their natal freshwater habitat to spawn. Because individuals are not in breeding condition every year, spawning migrations occur only once every 2-4 years. The arrow with the asterisk indicates that Arctic Char are iteroparous and can resume feeding migrations the following spring after spawning (3) Overwintering migrations (full grey arrows): in the years when individuals are not in breeding condition (resting (R)), they must still return to freshwater to overwinter to avoid lower temperatures and increased salinity in the marine environment. Because optimal conditions for spawning and overwintering may differ, use of non-natal overwintering sites might be more frequent than use of non-natal spawning sites. We hypothesized that individuals would favor less harsh migrations for overwintering, as indicated by the shorter river. Silhouette of adult fish from PhyloPic.org.demographic

A key variable driving the evolution of migrations and associated traits is migration distance. Migration distance greatly influences the costs of migrations, and strategies that minimize these costs will tend to be favored (Roff 2002). An example is the rapid evolution of a shorter migratory route in the warbler from continental Europe, the blackcap (*Sylvia atricapilla*), to take advantage of milder conditions in Britain (Berthold *et al*. 1992). Freshwater migration lengths and elevation gain (i.e. their ‘harshness’) are known to have a strong effect on the bioenergetic costs of migrations in anadromous fishes, including salmonids (Bernatchez & Dodson 1987). Migratory harshness can drive local adaptation of life history (Schaffer & Elson 1975), morphological (Crossin et al. 2004), and physiological traits (Eliason *et al*. 2011). While studies from Arctic Char have shown the influence of habitat accessibility on freshwater migrations (Gyselman 1994; Gilbert et al. 2016), the extent to which migration harshness drives habitat choice (particularly during overwintering migrations) and local adaptation remains largely unknown.

We combined double-digest RADseq (Andrews et al. 2016) and acoustic telemetry to study the migrations of anadromous Arctic Char from southern Victoria Island in the Canadian Arctic Archipelago (Fig. 2). The largest commercial fishery for Arctic Char in Canada has been operating in this region for over five decades (Day & Harris 2013) and the local Inuit people rely on this resource for economic, subsistence, and cultural purposes (Kristofferson & Berkes 2005). At least seven rivers in this region support populations of Arctic Char, five of which are commercially fished. Previous studies in the region have demonstrated that many of the stocks mix at sea (Dempson & Kristofferson 1987; Moore *et al.* 2016), and that genetic differentiation estimated from microsatellite markers is low (Harris *et al*. 2016). We first tested the hypothesis that Arctic char in the region home to natal rivers to spawn, but may overwinter in non-natal freshwater systems. This hypothesis predicts that individuals would use different freshwater systems in different years, but that this movement to non-natal streams would not necessarily result in gene flow. We used acoustic telemetry data to show that many individuals used different freshwater habitats in different years, and used population genomic data to infer the natal origins of migrating individuals, thus confirming that dispersal to non-natal habitats occurs. We used an Approximate Bayesian Computation (ABC) framework (Csilléry *et al.* 2010) to determine that an observed asymmetry in dispersal towards the Ekalluk River was not associated with increased gene flow in that direction. This further supports the hypothesis that dispersal does not necessarily lead to gene flow. We second hypothesized that overwintering migrations to non-natal freshwater systems would predominantly occur towards the system offering the least costly migration, here the Ekalluk River (Fig. 2; Table 1), given the central role of migratory harshness in driving survival and local adaptation in other salmonids. Consistent with our hypothesis, both the acoustic telemetry data and the population genomic data suggested a highly asymmetric pattern of dispersal towards the Ekalluk River. Finally, exploratory analyses using genome scans and genotype-environment associations identified outlier loci potentially associated with local adaptation. These markers were subsequently characterized using available genomic information from related salmonids. Together, our analyses provided new insights on patterns and consequences of migrations in a species of ecological, economic, and cultural importance.

**Figure 2.**
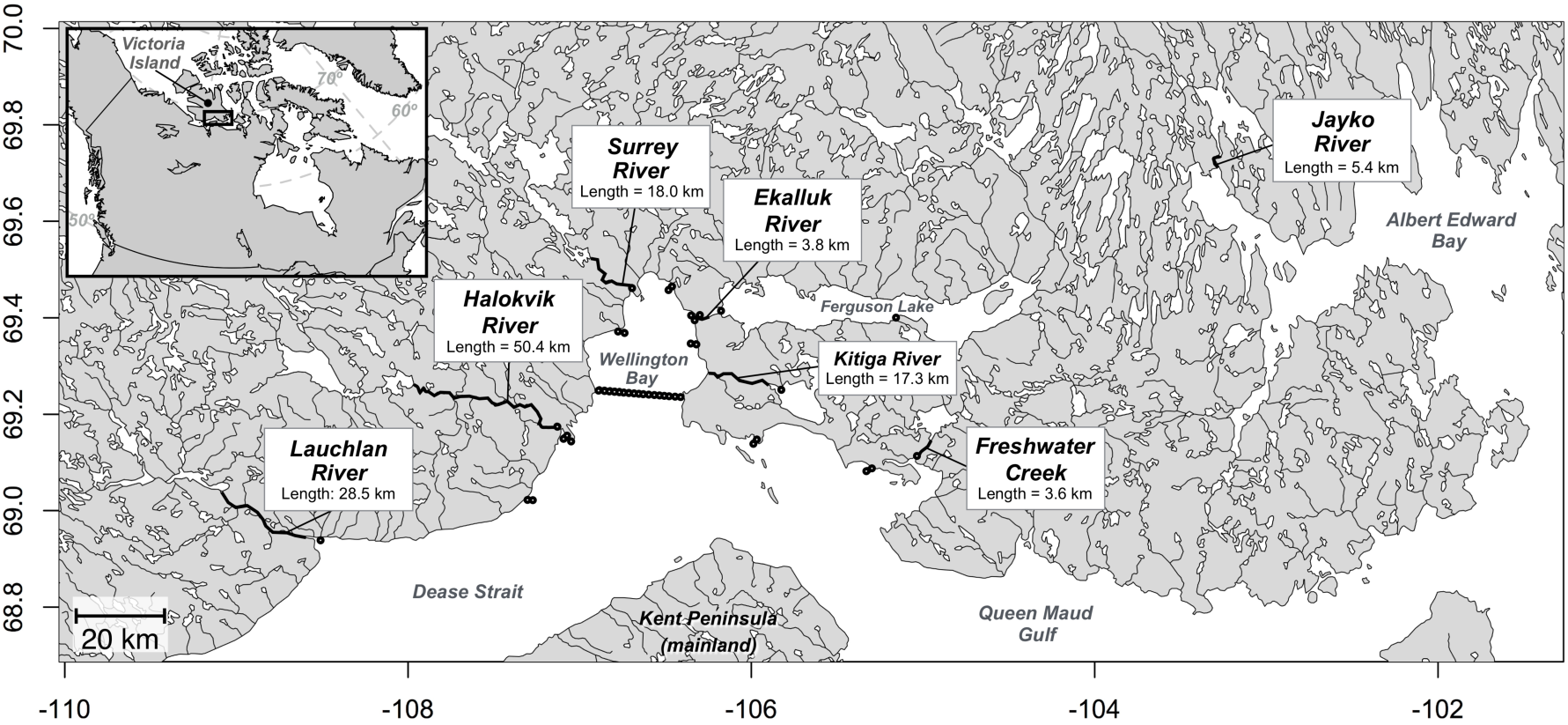
Map of the study area highlighting the names and lengths of the rivers sampled. The black circles with a white center represent the locations of the moored acoustic receivers used to track the movements of Arctic Char surgically implanted with transmitters at the Ekalluk, Surrey and Halokvik rivers.

**Table 1.** Sample information, river characteristics, and basic genetic diversity estimates for the six sampling locations used in the present study. Sample type refers to whether the individuals were collected specifically for DNA analysis (‘Baseline’) or were individuals that were equipped with an acoustic tag as part of the telemetry study (‘Tagged’). The sampling period refers to whether sampling targeted downstream migrating fish (‘Spring’) or upstream migrating fish (‘Fall’). *N*_*tot*_ is the total number of individuals sampled, *N*_<0.25 NAs_ the number of individuals with less than 25% missing data retained for all analyses, *H*_O_ the observed heterozygosity, *H*_S_ the expected heterozygosity, and *G*_IS_ is the inbreeding coefficient. All *G*_IS_ values are significant (<0.001).

| Sampling location | Coordinates | River length (km) | Elevation gain (m) | Sampling year | Sample type | Sampling period | $N_{tot} / N_{<0.25 \text{ NAs}}$ | $H_O$ | $H_S$ | $G_{IS}$ | $N_e$ (95% CI) |
| --- | --- | --- | --- | --- | --- | --- | --- | --- | --- | --- | --- |
| Lauchlan River | N68.94°<br>W108.52° | 28.5 | 64 | 2012 | Baseline | Fall | 57/57 | 0.102 | 0.105 | 0.03 | 335<br>(318 – 353) |
| Halokvik River | N69.16°<br>W107.09° | 50.4 | 118 | 2013 | Baseline | Fall | 60/59 | 0.099 | 0.103 | 0.037 | 823<br>(736 - 934) |
|  |  |  |  | 2013 | Tagged | Fall | 34/30 |  |  |  | - |
| Surrey River | N69.45°<br>W106.68° | 18 | 35 | 2013 | Baseline | Spring | 60/59 | 0.102 | 0.105 | 0.023 | 892<br>(798 – 1009) |
|  |  |  |  | 2014 | Tagged | Spring | 31/31 |  |  |  | - |
| Ekalluk River | N69.41°<br>W106.31° | 3.8 | 15 | 2013 | Baseline | Fall | 60/53 | 0.104 | 0.106 | 0.017 | 1081<br>(935 – 1280) |
|  |  |  |  | 2013 | Tagged | Spring | 30/26 |  |  |  | - |
|  |  |  |  | 2014 | Tagged | Spring | 31/31 |  |  |  | - |
| Kitiga River | N69.29°<br>W106.24° | 17.3 | 38 | NA | NA | NA | NA | NA | NA | NA | NA |
| Freshwater Creek | N69.12°<br>W105.00° | 3.6 | 7 | 1992 | Baseline | Unknown | 32/32 | 0.105 | 0.105 | 0.001 | 389<br>(351 – 436) |
| Jayko River | N69.72°<br>W103.28° | 5.4 | 3 | 2014 | Baseline | Fall | 60/58 | 0.104 | 0.107 | 0.022 | 368<br>(351 – 387) |

## Materials and Methods

### Telemetry data and sampling

Detailed methods for the acoustic telemetry data collection can be found in Moore *et al*. (2016). In short, 42 moored Vemco VR2W passive acoustic receivers were deployed between July 13 and September 8, 2013 (Fig. 2) along the southern shore of Victoria Island, Nunavut, near the community of Cambridge Bay. Detections were continually recorded until receivers were last retrieved between July 27 and August 31 2016. There are at least seven watersheds in the region that support runs of anadromous Arctic Char, and these watersheds are drained by rivers that vary in length and elevation gain (Fig. 2 & Table 1). We placed a receiver in six of these seven rivers to detect migrating fish (the Jayko River was excluded from the telemetry study because of the large distance between this site and the others). A total of 126 adult Arctic Char (>500 mm) were surgically implanted with Vemco V16 acoustic transmitters at three tagging locations (Table 1; only 124 were detected on the array and have telemetry data available). In all cases, the fish were collected at the river mouth near the ocean, and for the Ekalluk and Surrey River, tagging occurred during the downstream migration (early July), whereas tagging at the Halokvik River targeted upstream migrants (mid/late-August). A fin biopsy was taken from each tagged fish for genomic analysis. Additional baseline samples for genetic analysis were taken from sampling conducted during the upstream migrations to monitor commercial stocks or were collected from commercial fishery catches (Table 1). Samples from the Kitiga River (N69.29**°** W106.24**°**), which harbors an Arctic Char population, were also obtained, but DNA extractions for these samples failed. All of the sampling targeted adults but no information was available about the breeding status of sampled individuals. In the absence of this information, we refer to the use of non-natal freshwater systems as ‘dispersal’ throughout the manuscript, while we use the term ‘homing’ to refer to the use of natal habitat regardless of the purpose of the migration. The natal river of migratory adults cannot be determined with absolute confidence and is instead inferred from a combination of information on the sampling location, the telemetry data, and/or the genomic data. We also refer to all migrations back to freshwater in the fall as ‘return migrations’ regardless of whether an individual is dispersing or homing.

### Library preparation, sequencing, and SNP calling

A salt-extraction protocol adapted from Aljanabi & Martinez (1997) was used to extract genomic DNA. Sample quality and concentration were checked on 1% agarose gels and a NanoDrop 2000 spectrophotometer (Thermo Scientific). Concentration of DNA was normalized to 20 ng/µl (volume used = 10 µl; 200 ng total) based on PicoGreen assays (Fluoroskan Ascent FL, Thermo Labsystems). Libraries were constructed and sequenced on the Ion Torrent Proton platform following a double-digest RAD (restriction site-associated DNA sequencing; Andrews *et al.* 2016) protocol modified from Mascher *et al*. (2013). In short, genomic DNA was digested with two restriction enzymes (*PstI* and *MspI*) by incubating at 37^o^C for two hours followed by enzyme inactivation by incubation at 65^o^C for 20 minutes. Sequencing adaptors and a unique individual barcode were ligated to each sample using a ligation master mix including T4 ligase. The ligation reaction was completed at 22^o^C for 2 hours followed by 65^o^C for 20 minutes to inactivate the enzymes. Samples were pooled in multiplexes of 48 individuals, insuring that individuals from each sampling location were sequenced as part of at least 6 different multiplexes to avoid pool effects. One randomly chosen replicate per sampling location was also run on a different multiplex to evaluate sequencing errors (Mastreta-Yanes *et al.* 2015). Libraries were size-selected using a BluePippin prep (Sage Science), amplified by PCR and sequenced on the Ion Torrent Proton P1v2 chip. The obtained reads were aligned to the closely related Rainbow Trout (*Oncorhynchus mykiss*) genome (Berthelot *et al.* 2014) using GSNAP v2016-06-09 (Wu *et al.* 2016), then from these alignments STACKS v.1.40 (Catchen *et al.* 2013) was used for SNP identification, SNP filtering and genotyping (see online supplementary materials).

### Genomic data: basic statistics, population structure, and identification of putative dispersers

Observed (*H*_O_) and expected heterozygosity (*H*_E_), and the inbreeding coefficient (*G*_IS_) were estimated using GenoDive v2.0b27 (Meirmans and Van Tienderen 2004). Effective population size (*N*_E_) was estimated for the baseline samples using the linkage disequilibrium method in NeEstimator V2.01 (Do *et al.* 2013) with a critical value for rare alleles of 0.05. Pairwise population differentiation was quantified using *F*_ST_ (Weir & Cockerman 1984) and their significance values estimated with 1000 permutations also in GenoDive. A Discriminant Analysis of Principal Components (DAPC; Jombart *et al*. 2010) in the R package adegenet (Jombart 2008) was used to describe population structure. First, the ‘find.clusters’ function was used with the number of clusters *K* varying from 1 to 12. PCA scores of individuals were then plotted and the ‘compoplot’ function was used to calculate their proportion of membership to the genetic clusters identified. ADMIXTURE (Alexander *et al.* 2009) was also run varying the number of clusters *K* between 1 and 12. Putative dispersers were identified on the basis of cluster membership probabilities in DAPC and ADMIXTURE. Putative dispersers were defined as having a >75% probability of membership to a different cluster than the majority of other sampled individuals at that location. These putative dispersers, as well as individuals that had less than 75% probability of membership to any genetic clusters (i.e. putatively admixed individuals) were removed from some analyses (when noted) to avoid biases. Note that both of these thresholds are purposefully low to ensure all putative dispersers are removed.

### Population assignments

The software *gsi_sim* (Anderson *et al*. 2008) implemented in the R package *AssigneR* (Gosselin *et al*. 2016) was used to perform population assignments and test the power of assignment tests. We first simulated the assignment power of the baseline samples (see Table 1) and evaluated the impact of the number of markers used for assignment on power. Markers were ranked based on their *F*_ST_ values computed from a ‘training’ set of individuals comprising 50% of the total sample size per population and assignment success was computed for the remaining ‘hold-out’ individuals to avoid high-grading bias (Anderson 2010). The analysis was repeated for ten randomly generated training sets, and with ten randomly generated subsets of 30 individuals per population to eliminate any biases that could arise due to unequal sample sizes. We used the package *RandomForestSRC* (Ishwaran & Kogalur 2015) implemented in *AssigneR* to impute missing data based on allele frequencies per population using 100 random trees and 10 iterations and compared assignment success with and without imputations. We repeated these analyses on a dataset from which we removed the putative dispersers identified with the clustering analyses (see previous section for details). Next, we treated the acoustically tagged individuals as samples of unknown origin and assigned them back to the baseline samples using the function *assignment_mixture* in *AssigneR*. To avoid biasing the accuracy of the assignment with the missing data imputation, for each individual we only used the allele frequencies of the loci for which genotypes were present – the number of markers used for the assignments in this case therefore varied among individuals (min = 4,633; max = 5,966).

### Demographic simulations

To better understand the relative contributions of recent divergence and current gene flow on observed population structure, and to verify whether the observed patterns of dispersion translated into realized gene flow, an ABC pipeline using coalescent simulations was used (Csilléry et al. 2010). Following other approaches (e.g., Illera *et al*. 2014), three demographic models were compared and demographic parameters for the best model were estimated. The analyses focused on comparing the Ekalluk and Lauchlan samples and then the Ekalluk and Halokvik samples to limit computational demands. The datasets from which putative dispersers were removed were used for this analysis.

Furthermore, these comparisons are the most relevant to test our hypotheses as they are representative of the eastern (Ekalluk) and western (Lauchlan and Halokvik) group of populations identified as the first level of hierarchical structure (see Results). We tested three models: (1) a null hypothesis of panmixia; (2) a model of population divergence with gene flow (i.e. the isolation with migration model, IM); and (3) a model of population divergence without gene flow (strict isolation, SI). The simplest model of panmixia was controlled by a single parameter, *θ* = 4 *N*_*1*_µ, where *N*_*1*_ is the effective size of the panmictic population and µ is the mutation rate per generation. The SI model is characterized by six demographic parameters, namely *θ*1=4*N_1_µ*, *θ*_2_=4*N_2_µ*, and *θ*_A_=4*N_A_ µ*, with *N*_*1*_ and *N*_*2*_ the effective population size of the two daughter populations and *N*_*A*_ the effective size of the ancestral population, respectively. All these parameters are scaled by *θ*_1_=4*N_ref_µ* where *N*_*ref*_ is the effective size of a reference population. The two daughter populations diverged from the ancestral Tsplit generations ago with τ = T_split_/4*N_ref_.* The IM model was further characterized by the migration rates *M*_*1*_ = 4*N*_*m*_ (migration into population 1 from population 2) and *M*_*2*_ = 4*Nm* (migration into population 2 from population 1) sampled independently and where *Nm* is the effective migration rate. Large prior distributions following those commonly applied were used and drawn from uniform distributions (e.g., Sousa & Hey 2013). Coalescent simulations (n = 10^6^) were performed under each model using msnam (Ross-Ibarra *et al*. 2008), which is a modified version of the widely used ms simulator (Hudson 2002), under the infinite site model of DNA mutation. The pipeline of Roux *et al*. (2013) was used with modifications to test for panmixia, and priors were computed with a Python version of priorgen (Ross-Ibarra *et al*. 2008). Details of the summary statistics used, the model selection procedure, analysis of robustness, parameter estimations and posterior predictive checks can be found in the online supplementary materials. Values were not transformed into biological units since the mutation rate (µ) was unknown. Instead coalescent units were kept, and only ratios of *θ* and *M1/M2* were interpreted.

### Genome scans and functional annotation of outliers

Genome scans using different methods to identify outlier loci often only detect partially overlapping sets of loci (Gagnaire *et al*. 2015). A common practice to partially circumvent these problems is to combine genome scans with genetic-environment association (GEA) methods as a way to more reliably identify the most likely targets of selection (De Villemereuil *et al.* 2014). Therefore, we identified markers putatively under selection using two genome scan methods and a GEA method. First, Bayescan v1.2 (Foll & Gaggiotti 2008) was used to detect outlier loci with elevated *F*_ST_ among the baseline populations (Table 1) with 5,000 iterations, and a burn-in length of 100,000. To increase our chances of finding outliers while also limiting type-two errors, we tested values of 100, 1,000 and 5,000 for the prior odds (Lotterhos & Whitlock 2014). This analysis was run on the dataset from which putative dispersers were removed to avoid biases (n = 273 after dispersers removed). Second, we used PCAdapt (Luu *et al*. 2017), which uses PCA to control for population structure without *a priori* population definition, a particularly useful feature in the current study where we expect significant admixture among populations. Because of this feature, we ran the analysis on the full dataset including the putative dispersers, which also increased the analytical power associated with increased sample size (n = 318). Simulations have also shown that PCAdapt is less prone to type-2 errors than Bayescan, and to be less affected by the presence of admixed individuals in the samples (Luu *et al*. 2017). The optimal number of principal components necessary to describe population structure was determined using the graphical method described in Luu *et al.* (2017) and varying *K* from 1 to 10. Finally, we also used latent factor mixed models (LFMM; Frichot *et al*. 2013) implemented in the R package LEA to test directly for correlations between allele frequencies at specific loci and migratory difficulty. We also varied K from 1 to 10 to identify the optimal number of latent factors required to describe population structure. Migration harshness was measured as ‘work’ (Crossin *et al*. 2004), that is the product of river length and altitude gain, standardized so that the mean over all populations was of zero with a standard deviation of one. Because the environmental data had to be assigned to a sampling location, we used the dataset with the putative dispersers removed for this analysis (n = 273). As recommended by the authors in both user manuals, for PCAdapt and LFMM, the missing data were imputed as before using package *RandomForestSRC* (100 random trees and 10 iterations). All three approaches controlled for false discovery rates (FDR) with an *a* of 0.05.

An Arctic Char reference genome is in preparation (Macqueen et al. 2017), but it was not available at the time these analyses were conducted. Given the absence of a reference genome, we identified approximate locations for the markers by using the recently developed high-density linkage map for Arctic Char (Nugent *et al*. 2017). To increase the number of markers being positioned, the sex-averaged consensus map was used, including female- and male-specific markers. Although male salmonids have a low recombination rate in general (Sakamoto *et al.* 2000) and thus markers on the linkage map may not be as well positioned within linkage groups due to less frequent recombination, here we included them since we were interested in approximate general positions and the linkage group to which each marker belonged. Pairing of anonymous markers onto the linkage map was done using MapComp run iteratively, as previously described (Sutherland *et al.* 2016; Narum *et al.* 2017). In brief, MapComp aligns the flanking sequence of RAD markers of two genetic maps (here the Arctic Char genetic map and the anonymous Arctic Char markers) to a reference genome of a closely related species. As per Sutherland *et al.* (2016), here we used the Atlantic salmon genome as the closely related reference genome. Markers from each input comparison that are closest to each other in nucleotide position on the reference genome (and within a set distance) are then paired. The position of the paired marker is then given to the anonymous marker.

This pairing was conducted with a max distance of 1 Mb and done in 10 iterations, each time removing paired anonymous Arctic Char markers then rerunning the pairing with the Arctic Char genetic map to permit pairing of more than one anonymous marker with a single genetic map marker, as previously described (Narum *et al.* 2017).

All significant outliers were also used in a BLAST query against the Atlantic Salmon genome, which had the best annotation available at the time these analyses were conducted (Lien et al. 2016; NCBI Genome ICSASG_v2 reference Annotation Release 100; GCA_000233375.4). Outlier markers that were found by more than one method, but did not BLAST unambiguously to the Atlantic Salmon genome, were also checked against the Rainbow Trout (*Oncorhynchus mykiss*) genome (Berthelot *et al.* 2014).

## Results

### Telemetry data

The receivers located at the mouth of each river allowed the inference of likely freshwater overwintering/spawning sites of 98 acoustically tagged Arctic Char for up to three consecutive years (Table S1 in online supplementary materials). The telemetry data suggested a pattern of asymmetric dispersal, where many fish from all three tagging locations were detected one or more years in the Ekalluk River (Fig. 3). The Ekalluk River is the shortest and least harsh (i.e. least vertical and horizontal distance) river around Wellington Bay (Fig. 1 and Table 1), and one of the shortest in the entire area. Of the 47 fish tagged in both 2013 and 2014 at the Ekalluk River, 44 returned to this river each year (94%). Of the 21 fish tagged at the Surrey River, 16 returned to the Ekalluk River each year they were detected (76%), while only one fish returned to the Surrey River (5%). We therefore hypothesized that the fish tagged at the Surrey River were likely Ekalluk River fish that we intercepted moving north and west, an interpretation consistent with observations that many Ekalluk River fish visit the Surrey River estuary shortly after their out-migrations to the ocean (Moore *et al*. 2016) and also consistent with the population genomic data presented below. Of the 30 fish tagged at the Halokvik River, 28 returned to Halokvik River at least once (93%), and six returned to the Ekalluk River at least once (20%). Furthermore, of the 12 fish with more than one year of data that were inferred to have returned to the Halokvik River at least once, seven (58%) migrated to the Ekalluk River during other years, demonstrating that an individual might readily utilize different freshwater sites during its lifetime. In summary, a majority of fish from all three tagging locations (72%) migrating to the Ekalluk River at least once.

**Figure 3.**
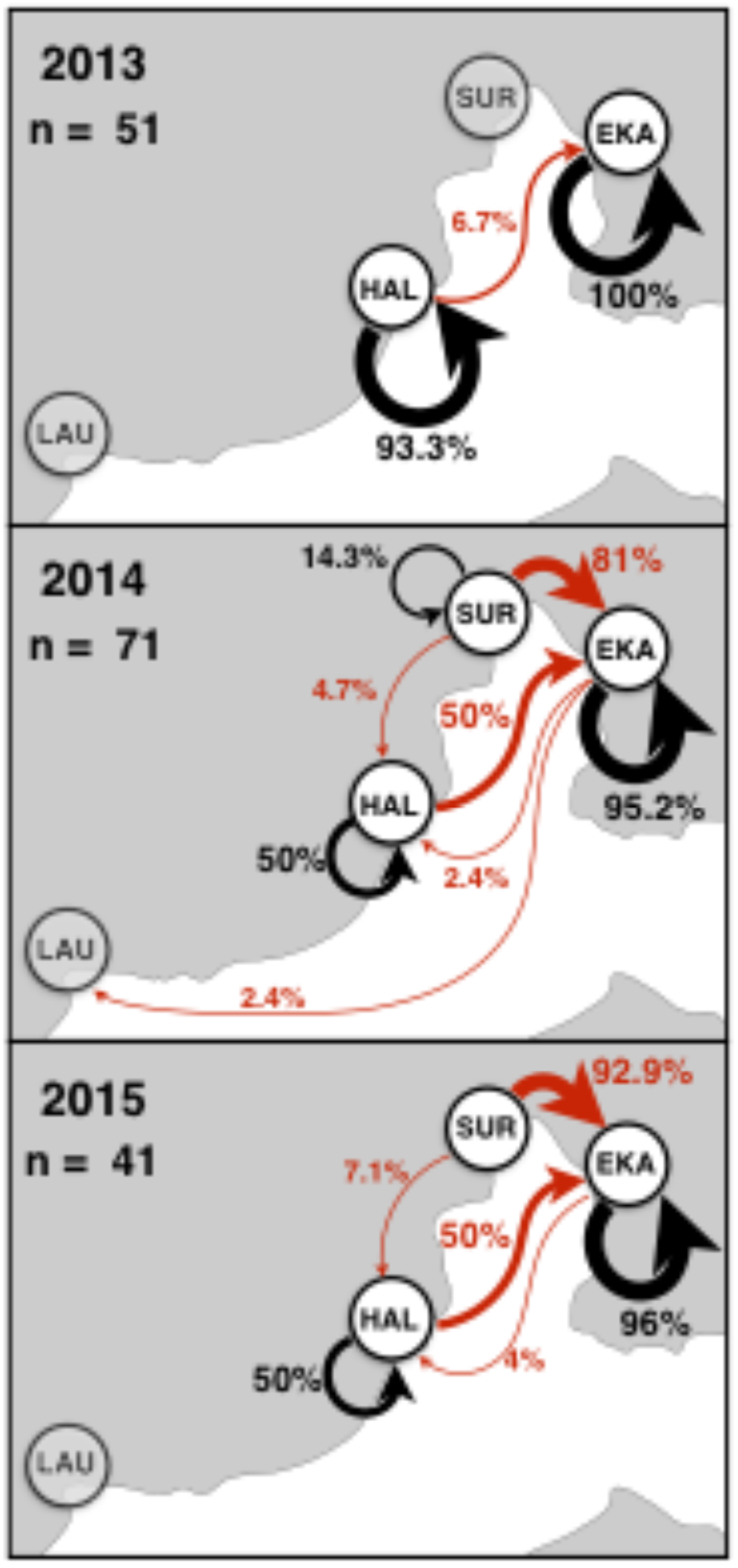
Choice of overwintering/spawning habitat by Arctic Char from three tagging locations inferred from acoustic telemetry data collected over three summers (2013, 2014, and 2015). The proportion of individuals returning to the same location where they were tagged (black) or dispersing to a different location from that where they were tagged (red) are indicated. The thickness of the arrows is roughly proportional to the proportion of individuals observed migrating to this location. The map indicates the approximate locations of Lauchlan (LAU), Halokvik (HAL), Surrey (SUR), and Ekalluk (EKA) rivers (shaded labels indicate no individuals were tagged there that year). The numbers of individuals that were unambiguously assigned to an overwintering location each year (n) are indicated.

### Sequencing and SNP calling

After cleaning and demultiplexing, a total of 1.9 billion reads were left with an average of 4.0 million reads per individual (coefficient of variation: 30.3%). The assembly resulted in a catalog containing 2,963,980 loci and a pre-filter total of 13,568,653 SNPs (over 1,272,427 polymorphic loci) after the population module of STACKS (Table S3 in supplementary materials). Individuals with more than 25% missing genotypes were removed from all analyses (Table 1), and high-quality SNPs were retained after filters (Table S3). In addition, 94 putatively sex-linked markers were removed (see Fig. S4 & S5 in supplementary materials and Benestan et al. 2017 for details) leaving a total of 6,136 SNPs used in all subsequent analyses (Table S3). Replicate individuals (n = 11) sequenced twice had identical genotypes at 92.0-97.1% of the markers (mean 94.6%), which was comparable to error rates reported in Mastreta-Yanes et al. (2015).

### Genomic data: basic statistics, population structure, and identification of dispersers

Levels of heterozygosity were comparable among populations (*H*_O_: 0.099-0.105; *H*_E_: 0.103-0.107), and although all *G*_IS_ values were significantly positive, indicating an excess of homozygotes, the values were all small (0.001-0.037) (Table 1). Effective population size estimates varied between 335 (Lauchlan River; 95% c.i.: 318-353) and 1081 (Ekalluk River; 95% c.i.: 935-1280). A weir enumeration study conducted in 1979-1983 found the Ekalluk River to be the most abundant population in the region, and the Lauchlan River the least abundant, but the estimates of census size were much larger than the *N*_E_ estimates reported here (183,203 for Ekalluk River and 10,850 for Lauchlan; McGowan 1990). Population differentiation among rivers was weak albeit significant, with an overall *F*_ST_ of 0.011 (95% c.i. 0.010-0.012; Fig 4a). The *F*_ST_ value between Ekalluk and Surrey rivers was substantially lower than the other values (*F*_ST_ = 0.001; 95% c.i. 0.0006-0.0016), which was in agreement with our telemetry observations that most fish captured at Surrey migrated to the Ekalluk River in the fall. It is thus likely that sampling at the Surrey River resulted in the interception of Ekalluk River fish migrating through the area, and those two sampling locations were combined for subsequent analyses unless otherwise noted. When the putative dispersers were removed (*see below*), the overall *F*_ST_ increased to 0.014 (95% c.i. 0.013-0.015; Fig. 4a). The presence of population structure between most sampling locations was also supported by the PCA analysis, although there was some important overlap among some sampling sites (Fig. 4b). The sampling locations that were most separated in the PCA (Jayko and Lauchlan rivers) were also the most geographically distant sampling locations. The PCA analysis was also repeated on a data set from which putative outlier loci were removed and returned essentially identical results (Fig. S8 online supplementary materials).

**Figure 4.**
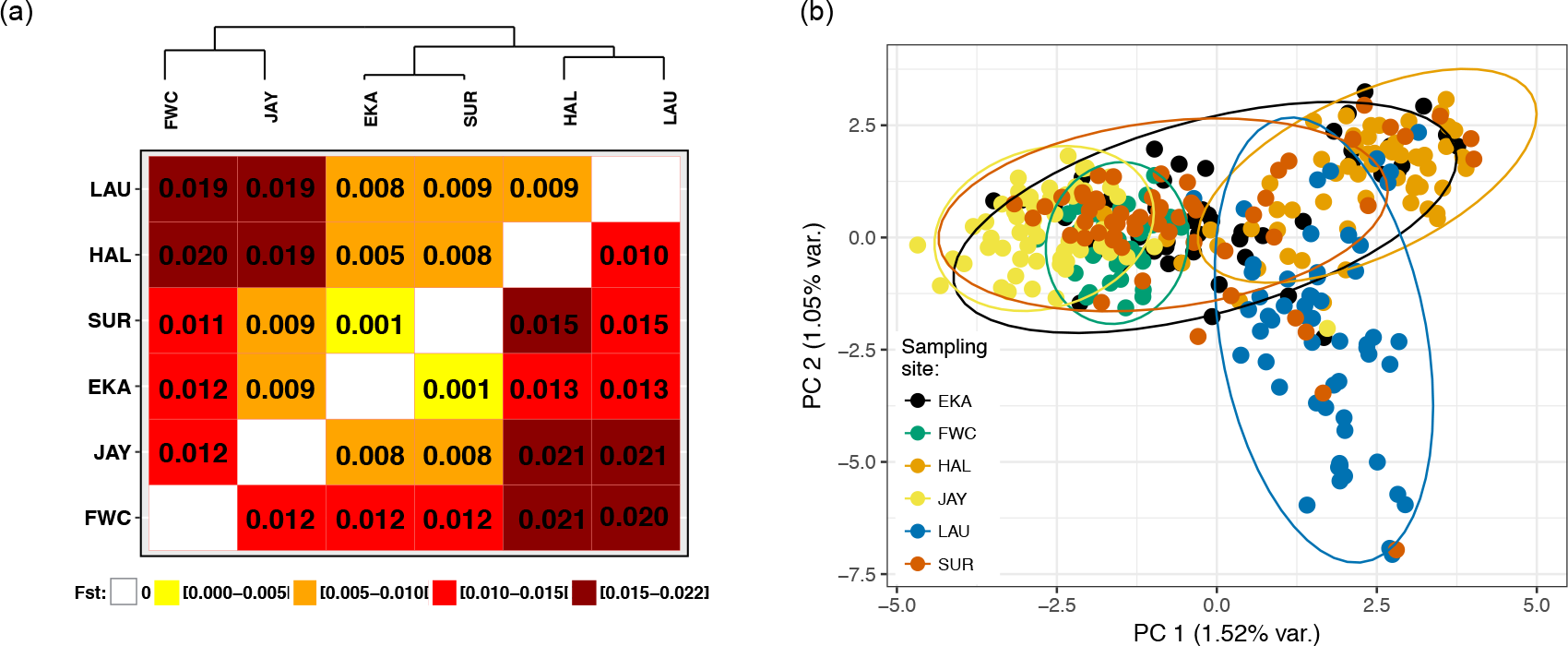
Description of population structure among the Arctic Char sampling sites: (a) heatmap of pairwise *F*_ST_ values before (above the diagonal) and after (below the diagonal) putative dispersers were removed from the samples (see text for details). All values are significantly different from zero (p <0.05). (b) Results of the principal components analysis showing PC scores of each individual along the first two principal axes (1.52% and 1.05% of the total variance explained respectively). Individuals and the 95% confidence ellipses are colour-coded by sampling location.

The Bayesian Information Criterion in the DAPC analysis best supported the presence of two genetic clusters (Fig. S6 online supplementary materials) differentiating the two westernmost populations (Lauchlan and Halokvik) from all others (Fig. 5). Based on cluster membership, several putative dispersers and admixed individuals could be identified (Fig. 5). A majority of putative dispersers (38/45; 85.4%) were individuals belonging to the western genetic cluster, but sampled in the Ekalluk or Surrey River. In contrast, many fewer dispersers from the eastern group were identified in the western sampling locations (three in Lauchlan and three in Halokvik; i.e., 13.3% of putative dispersers). In other words, the DAPC analysis supported the conclusion of asymmetric dispersal from the western sampling populations towards the shorter and less harsh Ekalluk River. Unlike the DAPC analysis, the cross-validation errors in the ADMIXTURE analysis did not support the presence of two genetic groupings (K = 1 had the lowest cross-validation error). Nonetheless, at *K* = 2, the individual Q-values were consistent with the results of the DAPC, and all putative dispersers identified with the DAPC had >50% probability of membership to the alternative genetic cluster in ADMIXTURE as well. Furthermore, at K=5 the ADMIXTURE results suggested that genetic differentiation among rivers was present, albeit weak.

**Figure 5.**
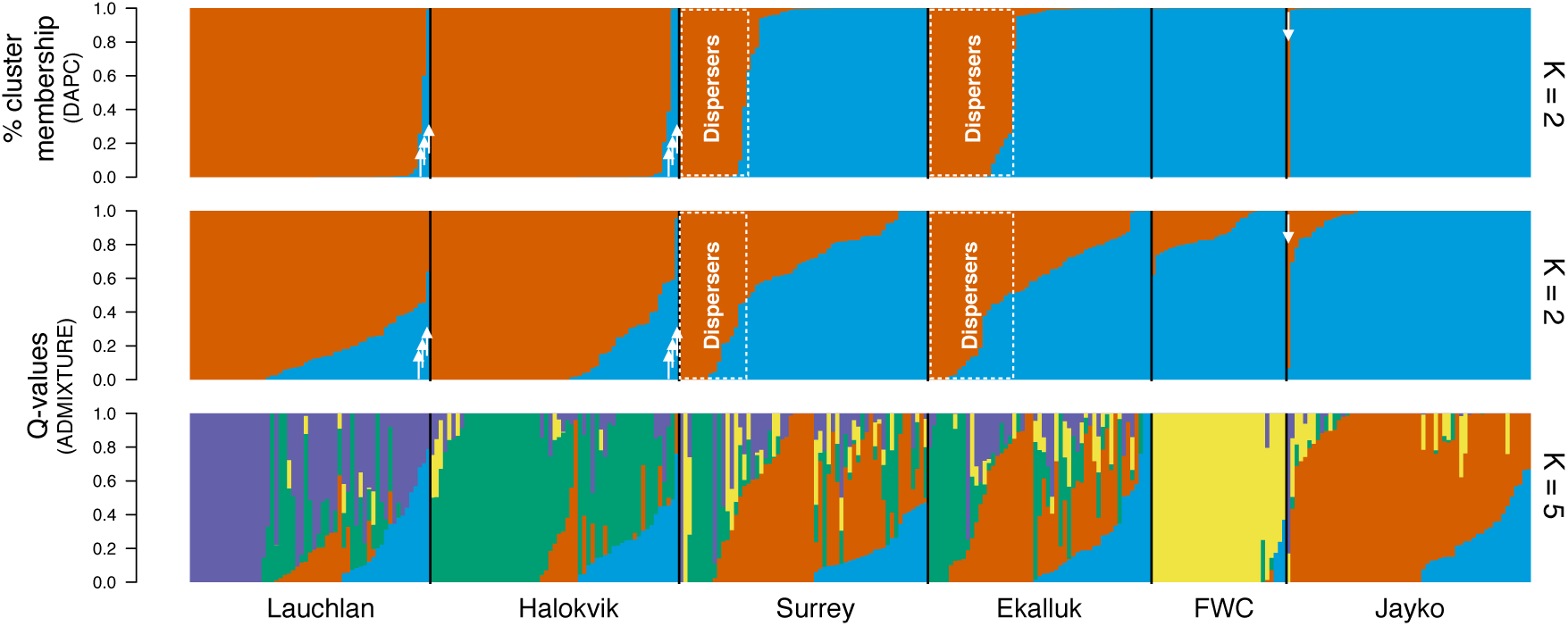
Results of two clustering analyses used to identify putative dispersers. The top panel is a compo-plot generated from a DAPC using two genetic clusters showing the percent cluster membership of each individual in the analysis. The bottom two panels are the individual Q-values from the ADMIXTURE analyses with two (middle) and five (bottom) clusters. The putative dispersers are individuals with Q-values or percent cluster membership greater than 0.75 but assigned to a cluster different from that of most other individuals sampled at the same sampling location and are identified with a white arrow or a white dotted rectangle. Note that the order of individuals varies among panels, but the identity of putative dispersers was the same in the two analyses.

### Population assignments

Overall, the results of the assignment tests performed on the baseline samples using all 6,136 markers revealed that there was sufficient power to infer the origin of individuals with high accuracy (lowest estimate including putative dispersers and without imputation: 83.1%; highest estimate excluding putative dispersers and with imputation: 92.7%; Fig. 6). When putative dispersers were included in the dataset, assignment success was lower overall (means over all sampling locations when all markers are included: 83.1% without imputations; 88.6% with imputations; range per sampling location: 66.0-100%; Fig 6a), and many individuals from Halokvik and Lauchlan River were assigned to Ekalluk and Surrey River, and vice-versa (Fig. 6b). As expected, assignment success was improved when the putative dispersers were removed from the dataset (means over all sampling locations when all markers are included: 88.2% without imputations; 92.7% with imputations; range per sampling location 78.5-100%; Fig. 6c), and most of the mis-assignments involved fish caught at Lauchlan River but assigned to Halokvik River, or fish caught at Jayko River assigned to Ekalluk and Surrey River (Fig. 6d).

**Figure 6.**
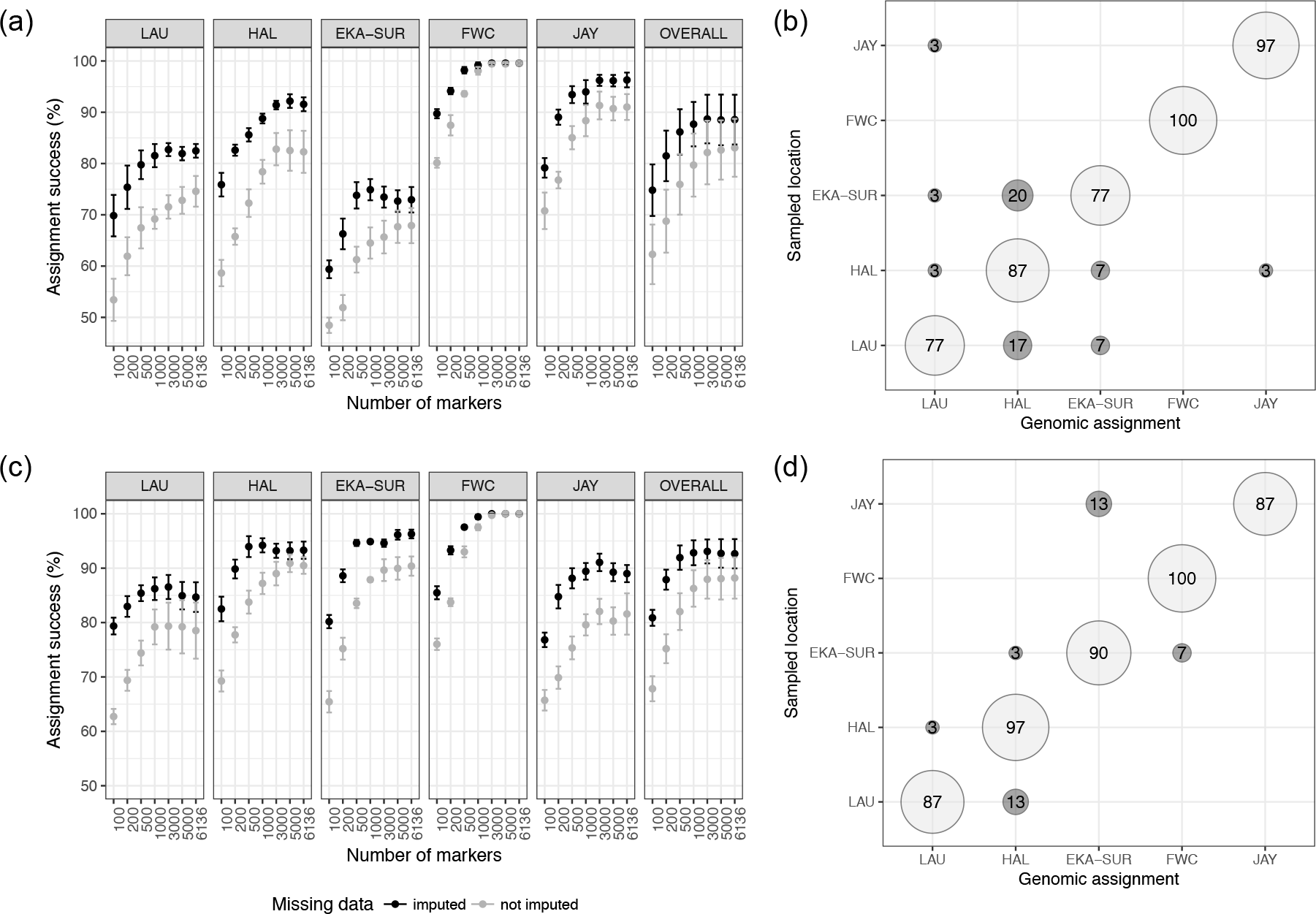
Results of assignment tests with all individuals (a & b) or with putative dispersers removed (c & d). The two left panels (a & c) show how assignment success per sampling location varied according to the number of markers used for the assignment, and on whether missing data were imputed using Random Forest (RF) or not. The two right panels (b & d) show results of the assignments with all markers with missing data imputed. The light grey circles on the diagonal display successful assignments, and the dark grey circles display mis-assignments (i.e., the sample was assigned to a different river from that where it was sampled). Circle size is proportional to the number of individuals in each category.

After confirming that there was sufficient assignment power, we used genomic data using all markers to assign the tagged individuals to their most likely river of origin (Table S1 online supplementary materials). Eleven of the 57 fish (19%) tagged at the Ekalluk River in 2013 and 2014 were assigned to the Halokvik River and two to the Lauchlan River (4%). Only two of the 11 putative Halokvik-origin fish were detected overwintering/spawning in the Halokvik River at least once, and all others were detected only in the Ekalluk system one or more years. In contrast, all but one (29/30) of the fish tagged at the Halokvik River (97%) were assigned to the Halokvik River itself. Six of those 29 individuals were detected overwintering/spawning in both Ekalluk and Halokvik River in different years. Notably, all but one fish (32/33) detected at least one year at Halokvik River were assigned to the Halokvik River. Finally, fish tagged at Surrey River were assigned to all possible rivers (except Freshwater Creek), but a majority (19/31) were assigned to the grouped Ekalluk-Surrey sample. In short, population assignments also support the prevalence of asymmetric dispersal towards the Ekalluk River.

### Demographic simulations

The model selection procedure unambiguously indicated that the isolation with migrations (IM) model was the best with P(IM) ~ 1, while both the panmixia and the strict isolation models had posterior probabilities close to zero (Table S5 in online supplementary materials). This inference was highly significant with a robustness of 1 (Fig. S9 in online supplementary materials). Estimates of demographic parameters from coalescent simulations produced confidence intervals of various widths, and differed most from the posterior in the Ekalluk-Halokvik comparison than in the Ekalluk-Lauchlan comparison. These estimates were mostly informative with regards to estimates of effective populations size, divergence times (except in the Ekalluk-Lauchlan comparisons), and intensity of migration rates, pointing especially to highly asymmetric gene flow (Fig. S10 & S11 in online supplementary materials). This asymmetric gene flow, however, was in the opposite direction of the asymmetric dispersal inferred from both the telemetry and the genetic data. Indeed, gene flow was higher from Ekalluk to Lauchlan, with a ratio of *M*_2_/*M*_1_ = 3.08, and from Ekalluk to Halokvik, with a ratio of *M*_2_/*M*_1_ = 4.06 (Table 2). Ratios of effective population size (θ_1_/θ_A_) tended to indicate an expansion of the Ekalluk population relative to the ancestral population in both comparisons (ratios of 9.68 and 3.45). Posterior distributions also indicated an expansion of the Halokvik population (θ_2_/θ_A_ = 2.58), but the Lauchlan population size was inferred to be smaller than the ancestral population size (θ_2_/θ_A_ = 0.082), although this should be interpreted cautiously since the estimate of the ancestral population size was not highly accurate. Posterior predictive checks in the Ekalluk-Lauchlan indicated that we were able to accurately reproduce the summary statistics. In the Ekalluk-Halokvik comparison, however, three statistics were significantly different from the observed data, namely the averaged number of shared polymorphic sites and the averaged and standard deviations of the net divergence (Da). In summary, the coalescent simulations and ABC approach provided robust estimates of demographic parameters, and suggested that the asymmetric dispersal towards Ekalluk River observed with both the telemetry data and the population assignment results did not translate into pronounced gene flow in that direction.

**Table 2.**
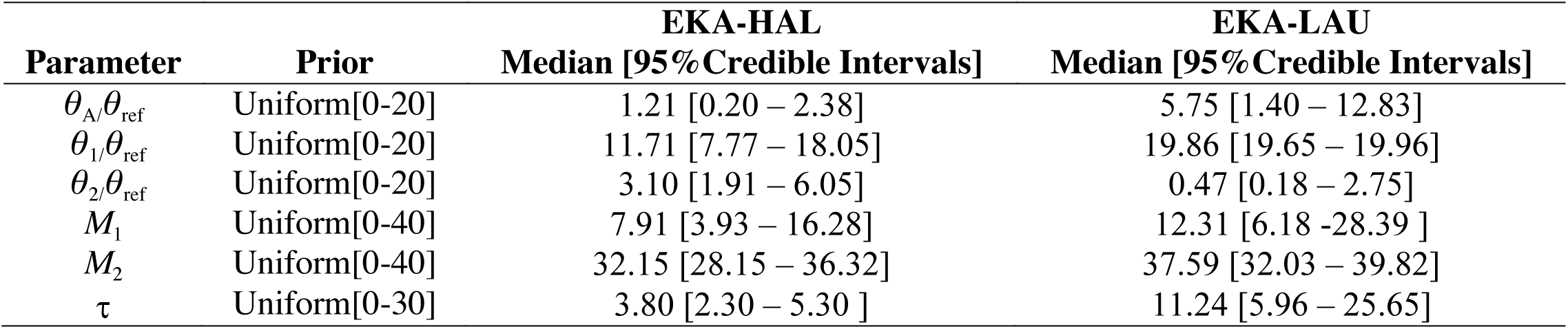
Parameter estimates for the isolation with migration (IM) model which was determined to have the highest posterior probability among the three models tested (P(IM)≈ 1) for two independent runs comparing Ekalluk and Halovik rivers (EKA-HAL) and Ekalluk and Lauchlan rivers (EKA-LAU). The parameters are *θ*A/*θ*ref for the ancestral population; *θ*1*/θ*ref and *θ*2/ *θ*ref for the two daughter populations (*θ*1 = Ekalluk in both cases and *θ*2 = Halokvik and Lauchlan depending on the analysis), with *θ*ref= 4*N*refµ where *Nref* is the effective size of a reference population and µ is the mutation rate per generation, *M*1 is the migration rate into population 1 (Ekalluk) from population 2 (Halokvik and Lanchlan) and vice-versa for *M*2, and τ = Tsplit/4*Nref* where Tsplit is the time (in generations) since population 1 and 2 diverged from the ancestral population.

| Parameter | Prior | EKA-HAL | EKA-LAU |
| --- | --- | --- | --- |
|  |  | Median [95% Credible Intervals] | Median [95% Credible Intervals] |
| $\theta_A/\theta_{\text{ref}}$ | Uniform[0-20] | 1.21 [0.20 – 2.38] | 5.75 [1.40 – 12.83] |
| $\theta_1/\theta_{\text{ref}}$ | Uniform[0-20] | 11.71 [7.77 – 18.05] | 19.86 [19.65 – 19.96] |
| $\theta_2/\theta_{\text{ref}}$ | Uniform[0-20] | 3.10 [1.91 – 6.05] | 0.47 [0.18 – 2.75] |
| $M_1$ | Uniform[0-40] | 7.91 [3.93 – 16.28] | 12.31 [6.18 -28.39 ] |
| $M_2$ | Uniform[0-40] | 32.15 [28.15 – 36.32] | 37.59 [32.03 – 39.82] |
| $\tau$ | Uniform[0-30] | 3.80 [2.30 – 5.30 ] | 11.24 [5.96 – 25.65] |

### Genome scans and functional annotation of outliers

The three different methods of outlier detection identified a total of 90 markers putatively under selection. The Bayescan analysis identified 30 outlier loci when the prior odds were set at 100 (Fig. S12 online supplementary materials), 10 loci with prior odds of 1000, and 5 loci with prior odds of 5000, all with an FDR of 0.05. The PCAdapt analysis was run assuming two genetic clusters after graphical evaluation of the eigenvalues according to Luu et al. (2017). Consistent with the DAPC results, the first two PC axes explained most variation and only those were retained to account for population structure (Fig. S13 online supplementary materials). Six outliers were identified with FDR = 0.05 (Fig S14 online supplementary materials). Of those six, four were also identified by the Bayescan run with prior odds of 100 and two with the run at prior odds of 5000. For the LFMM analysis, cross-entropy was also minimal at K = 2 (Fig. S15 online supplementary materials), and two latent factors were thus retained to control for population structure. With an FDR = 0.05, 58 markers were identified as outliers by the LFMM analysis, none of which overlapped with the markers identified by Bayescan or PCAdapt.

A total of 1405 of the 6136 markers were positioned onto the Arctic Char high-density linkage map (Nugent et al. 2017) (Fig. 7). Outliers were found on several different linkage groups throughout the genome. Of the 90 outlier markers, 34 provided unambiguous BLAST results against the Atlantic Salmon genome (see Table S6 for full list of successful BLAST results). Of those, 23 were found to be in a gene, while nine were found within 100kb of a gene. Only one of the four outliers that were significant in both the PCAdapt and the Bayescan had unambiguous BLAST results on the Atlantic Salmon genome, but three were located on the same 914Kb scaffold (*scaffold_363*) on the Rainbow Trout genome. The marker with the BLAST result was within the gene *FAM179B*, involved in the structure and function of primary cilia, an organelle usually associated with the reception of extracellular mechanical and chemical stimuli (Das *et al*. 2015). Interestingly, several genes close to outliers were involved in muscle or cardiac muscle function and development (*Myocyte-specific enhancer factor 2C*, *nebulette*, *glycogen phosphorylase muscle form*) or in gluconeogenesis (*fructose-1,6-bisphosphatase 1-like*), but a Gene Ontology analysis (*not shown*) failed to identify functions that were significantly enriched in the targets.

**Figure 7.**
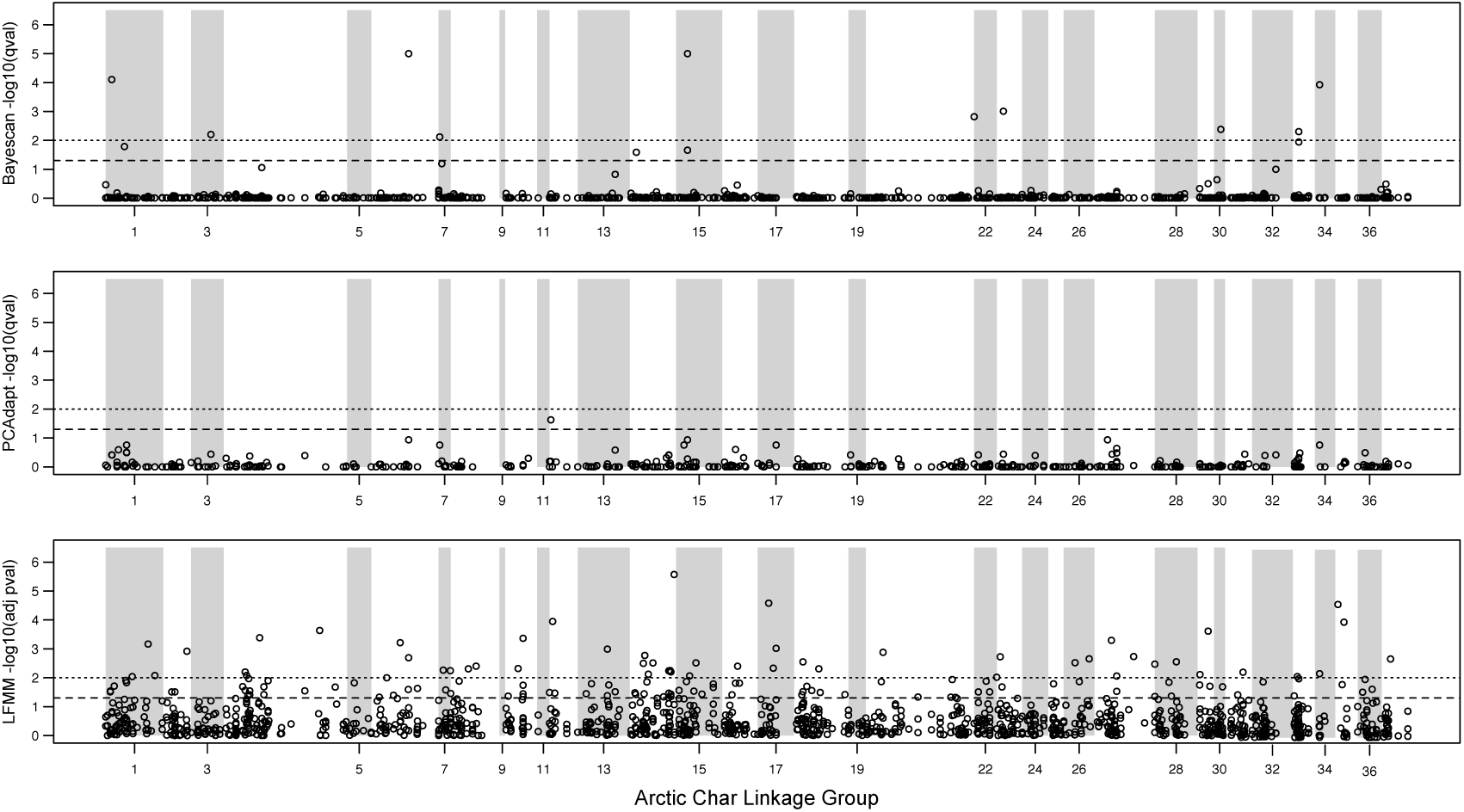
Manhattan plots showing the approximate locations of the 1405 markers successfully positioned on the Arctic Char genetic map along with multiple test corrected test statistics from two genome scan analyses (top: Bayescan and middle: PCAdapt) and a genetic environment association analysis correlating allele frequencies with migration harshness measured as ‘work’ (bottom: LFMM). Linkage groups are delineated by grey shading and x-axis labels. Dashed lines indicate 0.05 and 0.01 significance thresholds.

## Discussion

Concurrent developments in both genomic and telemetry technologies offer new and powerful ways to study the migratory ecology of animals behaving in their natural environments (e.g., Hess *et al*. 2014; Schaffer *et al*. 2016; Franchini *et al*. 2017). The present study illustrates how this integrative approach can be especially fruitful for animals displaying complex migratory behaviours and inhabiting logistically challenging regions. Here telemetry data suggested the presence of asymmetric dispersal between high Arctic populations of anadromous Arctic Char, and population genomic data helped to infer the river of origin of putative dispersers. Dispersal was predominantly from western sampling locations towards the Ekalluk River, the shortest and least harsh river in the Wellington Bay area. Coalescent simulations using an ABC approach also revealed an asymmetry in gene flow, but in the opposite direction, indicating that the observed asymmetric dispersal does not necessarily lead to gene flow. Instead, the coalescent simulations suggested that Ekalluk River, the most abundant Arctic char population in the region (McGowan 1990), was a source of gene flow to surrounding areas. Together, our observations suggested that Arctic Char home to their natal river to spawn, but may overwinter in rivers with the least harsh migratory route, thus potentially minimizing the costs of migrations in non-breeding years. Another finding facilitated by the integration of telemetry and genomic data was the identification of many outlier markers linked to genes demonstrated to have a role in muscle or cardiac function. Related physiological traits have been demonstrated to evolve in response to migration harshness in other anadromous salmonids (Eliason *et al*. 2011), thus providing a list of candidate genes that could be used as targets of future functional studies of migratory physiology in these Arctic Char populations.

### Asymmetric dispersal: overwintering habitat choice minimizes cost of migrations

Unlike most other anadromous salmonids, which return to freshwater exclusively to reproduce (Fleming 1998; Quinn 2005), Arctic Char and other *Salvelinus* species must return yearly to freshwater to overwinter (Armstrong 1974; Johnson 1980). Given that optimal habitats for reproduction and for overwintering likely differ, we might expect individuals to migrate to different locations depending on the purpose of the migration (Dingle 2014). Precise homing to reproduction sites is the norm in salmonids, and it has been suggested that local adaptation to nesting and rearing environments is an important driver of philopatry (Hendry *et al*. 2004). Requirements to home, however, might be relaxed when an individual is returning to freshwater to overwinter instead of to reproduce (i.e., when there are no genetic or evolutionary consequences). Accordingly, there is evidence that Arctic Char (and the closely related Dolly Varden; *Salvelinus malma*) home to their natal habitats to spawn, but can use non-natal habitats to overwinter (Armstrong 1974; Gyselman 1994; Moore *et al*. 2013; Gilbert *et al*. 2016). Here, we documented the use of alternative freshwater habitats by the same individuals in consecutive years using acoustic telemetry data. In addition, we used population genomic data to infer the natal origins of individuals, and confirmed that the asymmetric dispersal observed with the telemetry data was indeed due to fish of western origin (i.e., from Halokvik and Lauchlan) dispersing to the Ekalluk River. This was the case with the Ekalluk and Surrey baseline samples, which clustering analyses suggested contained a large proportion of western-origin fish. In addition, population assignment of the tagged fish showed that all but one fish detected at least once overwintering/spawning in Halokvik River were from Halokvik River, while many of the fish migrating to the Ekalluk River were from Halokvik River. The inference of asymmetric dispersal, therefore, is strengthened by the combination of independent and complementary sources of evidence from telemetry and genomic data.

The population genomic data could also be used to infer whether the observed dispersal resulted in high levels of gene flow. While dispersal was highly asymmetric towards Ekalluk River, the ABC modeling suggested that it did not necessarily translate into realized gene flow. Indeed, the best supported ABC model was the isolation with migration model with asymmetric gene flow, but the direction of the asymmetry was opposite to that observed with direct dispersal. This asymmetry in gene flow, instead, was from the most abundant population (Ekalluk) towards the least abundant populations (Halokvik and Lauchlan), as suggested by our *N*_E_ estimates (both from LDNE and ABC) and from field measures of abundance (McGowan 1990), and may thus simply reflect differences in effective gene flow. In summary, the observed asymmetry in dispersal does not translate into realized gene flow and suggests that dispersers from the Halokvik and Lauchlan rivers do not regularly reproduce in the Ekalluk River system.

Taken together, the observations of asymmetric dispersal with both genomic and telemetry data, but the lack of realized gene flow in the same direction, are consistent with the hypothesis that individuals in breeding condition home to spawn but may select overwintering habitats that minimize the costs of migrations in the years when they are not in breeding condition. Indeed, the Ekalluk River is one of the most easily accessible rivers in the region (measured as work; Table 1). While other rivers are also easily accessible (e.g., Freshwater Creek, Jayko River), they are further away from Wellington Bay, and comparatively fewer fish from the Wellington Bay tagging locations travel there (Moore *et al*. 2016). Given the lack of replication of this environmental axis in the current study, it is impossible to conclude without doubt that migration harshness is the main driver of overwintering habitat selection. Other features of the freshwater habitats may also explain the observed habitat choice. For example, the Ekalluk River drains the largest lake on Victoria Island, Ferguson Lake, which could offer more abundant or better overwintering habitats, perhaps resulting in lower overwintering mortality. Nonetheless, other studies of Arctic Char have also found spawning habitat accessibility to be a major constraint to migrations (Gyselman 1994; Gilbert *et al.* 2016), and the easier migrations afforded by the short Ekalluk River constitutes a plausible explanation for the observed patterns of dispersal. This hypothesis, however, assumes the ability of Arctic Char to assess which river provides the most readily accessible overwintering habitat. One possibility is that individuals explore a variety of freshwater habitats during their summer migrations and assess their accessibility. Observations of back and forth movement during homing in other salmonids (Quinn 2005), and the regular use of different estuaries by individuals during the summer documented by our telemetry work (Moore *et al.* 2016) suggest this is plausible. Another possibility is that the decision of where to overwinter relies in part on collective navigation (i.e., increased ability to navigate through social interactions), a possibility that has been discussed in relation to salmon homing (Berdahl *et al*. 2014). Given that the Ekalluk River has the most abundant population of Arctic Char in the region (McGowan 1990; Day & Harris 2013), it is also possible that collective navigation biases dispersal towards the river with the greater number of individuals. Continued collection of telemetry data will allow future tests of some of these hypotheses and may help in specifically testing the idea suggested by our observations that migration harshness drives overwintering habitat choice.

In summary, the integration of telemetry and genomic data allowed nuanced insights into the complex migratory ecology of anadromous Arctic Char. Notably, it allowed circumventing logistical challenges posed by the sampling of baseline samples. In most salmonids, sampling in the freshwater is used to ensure the origin of sampled fish for baseline collections, but here the use of different freshwater habitats by adults within their lifetime precludes any certainty in inferring origins of sampled adults even if collected in freshwater. Sampling juveniles prior to their first marine migration (e.g., Moore *et al*. 2013) would also be an option, but attempts at collecting juveniles in 2010 and 2015 failed: seine nets, minnow traps, and electrofishing were unsuccessful, (the latter possibly affected by the low conductivity of Arctic waters), the watersheds under study are vast and inaccessible, and the location of juvenile rearing habitats are unknown. Given the complex migratory behaviour of Arctic Char and the logistical challenges associated with sampling, the integration of telemetry and genomic data provided a powerful tool for understanding the ecology of this species.

### Evidence for local adaptation in migratory traits

Anadromous salmonids display tremendous diversity in migratory traits both among and within species, and their study has contributed to our understanding of migration ecology (Hendry *et al*. 2004; Dingle 2014). The application of molecular tools has provided important insights to the study of the genetic basis of migratory traits in salmonids. For example, O’Malley & Banks (2008) used a candidate gene approach to link variation in two circadian rhythm genes, *OtsClock1a* and *OtsClock1b*, to variation in run timing in anadromous Chinook salmon (*O. tshawytscha*). Other studies have identified SNP markers linked to migration timing in both Chinook salmon (Brieuc *et al.* 2015) and steelhead trout (*O. mykiss*; Hess *et al*. 2016). Finally, Hecht *et al*. (2015) linked variation in RAD markers with 24 environmental correlates using redundancy analysis in Chinook salmon, finding that freshwater migration distance was the environmental variable explaining most of the genomic variation. This last result parallels our own observations suggesting the importance of migratory harshness, and is also corroborated by many classic studies demonstrating the importance of migratory difficulty as a selective factor driving adaptation in morphological, physiological, and life-history traits (Shaffer & Elson 1975; Bernatchez & Dodson 1987; Crossin *et al*. 2004; Eliason *et al*. 2011).

In the present study, we used genome scans (Bayescan and PCAdapt) and genetic-environment correlations (LFMM) with migratory harshness to identify markers putatively under divergent selection. These approaches have been used extensively to identify putative targets of selection, but also have several limitations, which have been amply discussed in the literature (Lotterhos & Whitlock 2014; Haasl & Payseur 2015; Bernatchez 2016; Hoban *et al*. 2016). For example, recent demographic history and patterns of isolation by distance have been shown to lead to high levels of false positives (Lotterhos & Whitlock 2014; Hoban *et al*. 2016). The geographical proximity of the populations under study, however, precludes important differences in terms of demographic history since the re-colonization of the area from a single source population in the last 6,500 years (Moore *et al*. 2015). In addition, Lotterhos & Whitlock (2015) concluded that maximum power in genome scan studies could be achieved by examining geographically proximate and genetically similar populations that experience contrasting environments, criteria that our system fulfilled. Nevertheless, few of the outlier markers identified by the three methods overlapped, a common finding (de Villemereuil *et al*. 2014; Gagnaire *et al*. 2015).

In accordance with our *a priori* expectation that migration harshness might be a selective factor differing among rivers, several of the outlier SNPs were linked to muscle and cardiac functions and development. One of the outlier SNPs was located within 75 kb (5’ side) of *myocyte enhancer factor 2* (MEF2), a transcription factor acting as an important regulator of vertebrate skeletal muscle differentiation and heart development (Potthof & Olson 2007). Supporting the relevance of this protein for migrating salmonids, cardiac mRNA levels of MEF2 were elevated in Atlantic salmon following 10 weeks of experimental exercise training (Castro *et al*. 2013). An outlier was also located within the gene coding for *nebulette*, a cardiac-specific actin-binding protein essential in the structure of the sarcomere Z-disc (Bonzo *et al*. 2008). Another marker was within the gene coding for *muscle glycogen phosphorylase* (PYGM), the enzyme responsible for breaking up glycogen into glucose subunits to power muscle cells (Kitaoka 2014). Finally, another marker was within the gene coding for *fructose-1,6-biphosphatase*, which is involved in the conversion of glycerol into glucose (i.e., gluconeogenesis; Lamont *et al*. 2006). The abundance of putative targets of selection identified linked to muscle and cardiac functions and development is interesting in the context of variation in migration harshness. A Gene Ontology (GO) analysis, however, did not identify any functions that were significantly enriched. Nevertheless, the list of outliers identified provides interesting candidates for future functional studies on the physiology of migrating Arctic char that might help solidify the causal link between the migratory environment and genomic variation. This work will be further aided by the upcoming availability of an Arctic Char reference genome (Macqueen *et al*. 2017).

### Conclusions

Here we used an integrative approach to link genome-wide data with telemetry and concluded that extensive dispersal to overwintering habitats is not necessarily associated with high gene flow. These movements were biased towards a short river suggesting that migration harshness might drive overwintering habitat choice in anadromous Arctic Char. Genome scans for outliers and genetic-environment association analysis identified several markers in genes associated with muscle and cardiac functions, further suggesting the importance of migratory harshness in driving local adaptation. Future work in this system will build on these results to describe physiological differences among populations in relation to migration harshness. These will be particularly relevant in the context of adaptation to a changing Arctic, where migratory environments will probably exert an important selective pressure on populations (e.g., Eliason *et al*. 2011). Many of our conclusions would have been difficult, or impossible, to reach with genomic or telemetry data alone, and our study therefore illustrated the synergies made possible by combining the two types of data (Shafer *et al*. 2016). Such integrative approaches will continue to increase our understanding of how migratory behaviour interacts with gene flow to influence the spatial scale at which local adaptation can evolve, and will advance the study of the genetic basis of migratory traits of species behaving in their wild habitats.

## Acknowledgements

We are very grateful for the support of the Ekaluktutiak Hunters and Trappers Organization and of the residents of Cambridge Bay, who made this work possible. Koana! We also acknowledge Kitikmeot Food Ltd. for logistical support and their precious collaboration with the plant sampling program. We thank B. Boyle and G. Légaré at the IBIS sequencing platform for their help and advice regarding library preparation and sequencing. A.-L. Ferchaud, L. Benestan, T. Gosselin, S. Bernatchez, A. Perreault-Payette, & C. Perrier shared scripts and provided advice on data analysis and interpretation. The telemetry work was supported by the Ocean Tracking Network (OTN) through a network project grant (NETGP No. 375118-08) from Natural Sciences and Engineering Research Council of Canada (NSERC) with additional support from the Canadian Foundation for Innovation (CFI, Project No. 13011), by the Polar Continental Shelf Project of Natural Resources Canada (grant #107-15), and by in kind logistical support from the Arctic Research Foundation. J.-S. Moore was supported by fellowships from the Fonds Québécois de Recherche sur la Nature et les Technologies and the W. Garfield Weston Foundation. J. Le Luyer, B. Sutherland and Q. Rougemont were supported from various grants from the Canada Research Chair in Genomics and Conservation of Aquatic Resources led by L. Bernatchez.

## Author contribution

J.-S.M. and L.N.H. designed this study and conducted the fieldwork. J-SM performed part of the lab work, most of the analyses, and wrote the manuscript. J.L. identified the SNPs using the STACKS workflow. B.J.S. mapped the markers using MapComp and contributed to the functional annotations. Q.R. performed the ABC analyses. R.F.T., A.T.F., and L.B. provided guidance and supervision at various stages of the study. All authors contributed significantly to the writing of the article.

## Data accessibility

The .vcf files, the R code, the sequences of the RAD tags and the results of the genome scans and GEA analyses used in the construction fo the Manhattan plot can be found on Dryad (<u>http://dx.doi.org/10.5061/dryad.f3sm9.2</u>). The raw reads will be deposited to NCBI Short Read Archive. The pipeline used for the ABC simulations will be available at https://github.com/QuentinRougemont/abc_inferences. The pipeline for collecting and preparing the Arctic Char genetic map, running MapComp iteratively, and plotting the results are available at: https://github.com/bensutherland/salp_anon_to_salp

